# An expanded microsporidia species attribute database documents the extensive ecological and phenotypic diversity of these parasites

**DOI:** 10.64898/2026.09.21.753277

**Authors:** Carolyn Chen, Amel Alassal, Aaron W. Reinke

**Author notes:** These authors contributed equally (Carolyn Chen, Amel Alassal).

## Abstract

Microsporidia are a group of parasites that exhibit vast morphological, ecological, and genetic diversity. To catalog this diversity, we previously generated a database of attributes for microsporidia species. Here we expand this database to include species described since 2021, previously missed descriptions, and provisional species defined by clustering GenBank 18S rRNA sequences, resulting in 1,795 species. Using these data, we describe when and where microsporidia have been reported, their hosts and host habitats, the tissues they infect, and their spore morphology. A phylogeny of 18S sequence clusters shows that closely related microsporidia tend to infect closely related hosts, such that sequence identity is predictive of host taxonomy. Clustering also reveals previously unappreciated genetic diversity within some named species, in particular the human pathogen *Vittaforma corneae*, suggesting that these may comprise multiple species. We have also generated a web portal through which new species information can be submitted, which will facilitate regular updates. This database provides a resource for systematic analysis of microsporidia properties and a reference for taxonomic descriptions.

**Author summary:** Microsporidia are parasites that infect animals from insects and fish to humans, and species vary enormously in the hosts they infect, the environments they occupy, and the appearance of their spores. This information has accumulated over more than 150 years of species descriptions scattered across thousands of publications. Here we expand a database that brings it together, adding species described in the last five years, older descriptions that had been missed, and species known only from DNA sequences in public repositories. The resulting database of nearly 1,800 species allows us to show when and where microsporidia have been discovered, which animals and tissues they infect, and how their spores vary in size and shape. Comparing sequences across species shows that closely related microsporidia tend to infect closely related hosts, so a parasite’s DNA alone suggests the kind of animal it infects. It also reveals that some described species, most notably the human eye pathogen *Vittaforma corneae*, contain far more genetic diversity than their names suggest and likely represent several distinct species. We also provide a web portal through which researchers can submit new species, providing a lasting resource for studying and classifying these parasites.

## Introduction

Microsporidia are a group of obligate intracellular fungal parasites that infect hosts throughout the animal kingdom as well as many protists [1,2]. More than 1,600 species in over 200 genera have been described, and these species display remarkable diversity in nearly every property that has been examined [3]. Individual species range from specialists of a single host to generalists infecting hosts from multiple phyla, and collectively microsporidia occupy terrestrial, freshwater, marine, and brackish environments [4]. Infection can be restricted to a single tissue or spread systemically, and the infectious spore, defined by its unique polar tube, varies widely between species in size, shape, and polar tube length [5]. This diversity has practical consequences as microsporidia are opportunistic pathogens of humans and cause economically important diseases of honey bees, silkworms, and farmed shrimp [6–9]. Microsporidia can also be beneficial, with some species suppressing malaria parasites in mosquitoes and others serving as biological control agents of insect pests [10,11].

Knowledge of microsporidia diversity is dispersed across more than 150 years of species descriptions [12]. These descriptions typically report a common set of attributes, including the host and site of infection, geographic locality, spore morphology and dimensions, and the number of polar tube coils. To make this information accessible for systematic analysis, we previously assembled these attributes into a database covering ∼1,440 microsporidia species entries [4].

In parallel, sequencing has become central to cataloging microsporidia; the 18S rRNA gene is the most commonly sequenced marker and is often the only molecular data reported for a new species [3]. Together with genome-scale phylogenies, 18S trees define the current clade-level classification of the phylum, and 18S sequence diversity has been used to delineate species [3,13,14]. Because 18S can be amplified directly from hosts and environmental samples, it is also widely used to survey microsporidia diversity independent of formal species descriptions [15,16]. GenBank now contains thousands of microsporidia 18S sequences from such surveys, representing a growing body of diversity absent from attribute-based catalogs.

Here we present an expanded microsporidia species attribute database. We curated species described in the last five years, used large language model-assisted searches to recover previously missed descriptions, and clustered GenBank 18S rRNA sequences into species-level units, resulting in a database of 1,795 species. We determined the taxonomy and habitat of each reported host and used these data to describe when and where microsporidia species have been found, the hosts and tissues they infect, and their spore morphology. We also constructed an 18S phylogeny of species-level clusters, which relates host specificity and habitat to phylogenetic relatedness and reveals unappreciated genetic diversity within described species. Finally, we generated a web portal that allows community submission of species information to support continued updates of this resource.

## Results

### Generation of an expanded microsporidia attribute database

Our previously described database contained 1,440 entries, each recording attributes of a single microsporidium, including its hosts, locality, sites of infection, spore morphology, and associated sequence accessions [4]. We first consolidated 16 redundant records, retaining 1,424 species (S1 Fig). We then added species reported from 2021 to 2026, identified through targeted literature searches, which contributed 64 species, and performed broader LLM-assisted searches of the literature, which identified 80 additional species that had been missed previously (S1 Fig).

To capture diversity known only from sequencing, we retrieved all microsporidia 18S rRNA sequences from GenBank, trimmed them to a common region using a microsporidian covariance model, and clustered them into species-level units (S1 File). To select a clustering threshold, we evaluated identity cutoffs from 95% to 100% against species labels from sequence metadata and chose 98% identity, which maximized agreement between clusters and binomial species names (S2 Fig). Each cluster was matched to the database by shared accession or species name. Clusters without a database match were added as provisional species with unique identifiers, provided at least one of their sequences had an annotated host, contributing 227 sequence-defined species. Named species were represented by clusters of 1 to 705 sequences, 53% of which contained more than one sequence, whereas provisional species were represented by clusters of 1 to 29 sequences, 33% of which contained more than one sequence. The final database contains 1,795 species, comprising 1,245 named and 550 provisional species, with one third of species (n = 606) represented by at least one 18S accession (S1 Fig, S1 Table). Each 18S cluster with its constituent accessions and associated host, locality, and habitat records is listed in S2 Table.

### Microsporidia display extensive diversity across many dimensions

The expanded microsporidia attribute database allows the depiction of microsporidia diversity. We first examined when microsporidia species have been reported. Descriptions span from the 1850s to the present, with the rate of discovery increasing sharply from the 1960s and exceeding 100 species per decade (Fig 1A). The proportion of provisional species has grown in recent decades, reflecting the increasing contribution of sequence-based reports. Microsporidia have been reported from countries on every continent, except Antarctica, as well as from open ocean, though records are concentrated in North America, Europe, Russia, and parts of Asia, likely reflecting where microsporidia discovery efforts have been focused, rather than the true distribution of these parasites (Fig 1B).

**Fig 1.**
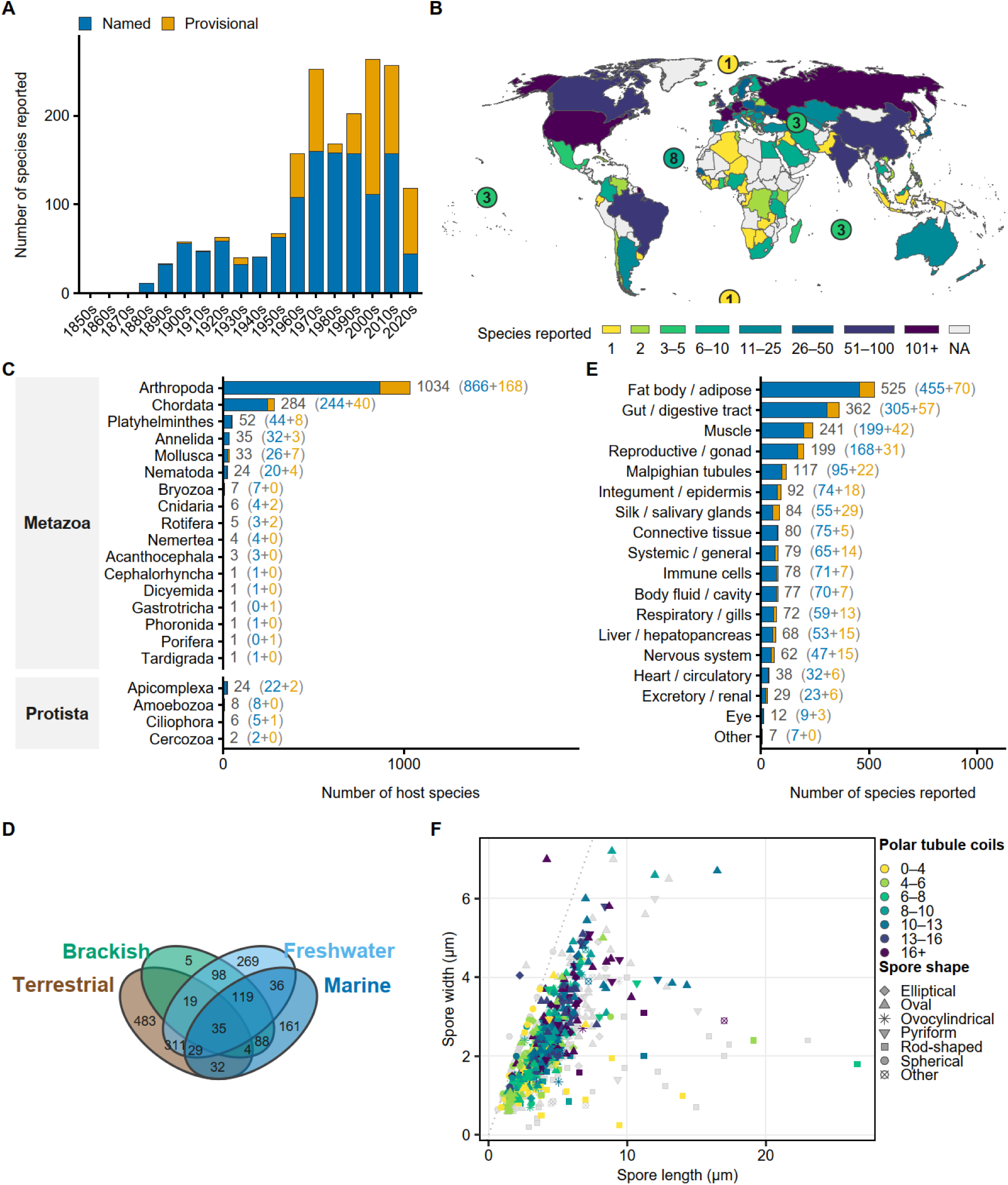
Temporal, geographic, host, habitat, tissue, and morphological diversity of microsporidia species. **(A)** Number of microsporidia species reported per decade, coloured by whether the species has a formal binomial (named) or not (provisional) (*n* = 1,776 species). **(B)** Geographical location of microsporidia species, with each species depicted once for each country where it has been reported. The number of species in each country is depicted as a heat map according to the scale below. Numbers in circles describe species found in respective oceans (*n* = 1,613 species). **(C)** Number of host species reported to be infected, by phylum and grouped by kingdom. Hosts infected by at least one named microsporidia species are shown in blue, and hosts infected only by provisional species are shown in orange. Values in parentheses give the named and provisional components of each total (*n* = 1,533 host species). **(D)** Venn diagram of the major habitats inhabited by the hosts of each microsporidia species (*n* = 1,689) microsporidia species. **(E)** Number of microsporidia species reported to infect each tissue type (*n* = 1,202 species) coloured as in panel A. **(F)** Spore length and width dimensions, with spore shape indicated by symbol and the number of polar tubule coils indicated by colour according to the legend at the right (*n* = 940 species). Species without reported coil numbers are shown in grey.

To characterize the hosts of microsporidia, we extracted each reported host and determined its taxonomy by querying four taxonomic databases (S3 Table). The 1,533 host species span 17 metazoan and four protist phyla (Fig 1C). Arthropods are by far the most commonly reported hosts (1,034 species), followed by chordates (284), with smaller numbers of platyhelminth, annelid, mollusc, and nematode hosts. Microsporidia also infect other parasites, including apicomplexan gregarines that themselves infect animal hosts. The hosts of provisional species are also distributed across this same taxonomic breadth. Since the last update of the database, the one new phylum is Tardigrada.

We next determined the habitat of each host using habitat annotations from these same taxonomic databases, supplemented with a curated table of genus-level habitats for hosts these databases did not resolve; assignments from the two approaches were largely concordant (S3 and S4 Tables, S3 Fig). The habitat of each microsporidia species was taken as the union of its hosts’ environments. Terrestrial and freshwater environments account for the majority of species, with 483 species having exclusively terrestrial hosts, 269 exclusively freshwater, 161 exclusively marine, and 5 exclusively brackish (Fig 1D). A substantial fraction of species span multiple habitats, most commonly terrestrial and freshwater.

Microsporidia infect nearly every host tissue. Sorting reported infection sites into 17 categories, the most commonly infected tissues are adipose tissue, the digestive tract, muscle, and reproductive tissue, with many species reported to infect multiple tissues and 79 described as systemic or generalized infections (Fig 1E).

Spore morphology is similarly varied. Reported spores are most commonly oval or pyriform, with lengths ranging from ∼1 to over 25 μm (Fig 1F). The number of polar tube coils ranges from fewer than four to more than 16, with larger spores tending to have more coils.

### Microsporidia phylogenetic tree displays diversity and relationships between host specificity and phylogenetic relatedness

To relate these attributes to evolutionary relationships, we built a maximum-likelihood phylogeny from representative 18S sequences from each species and displayed the 609 species-level clusters with at least one reported host, annotated with species status, clade assignment, host habitats, and host taxonomy (Fig 2, S4 Fig). The tree recovers the major recognized microsporidia clades, including Metchnikovellida, Ovavesiculida, Glugeida, Nosematida, Enterocytozoonida, Amblyosporida, and Neopereziida, and clade assignments agree with a previous assessment for all overlapping species, though the minor clade Caudosporida is not recovered as monophyletic [3].

**Fig 2.**
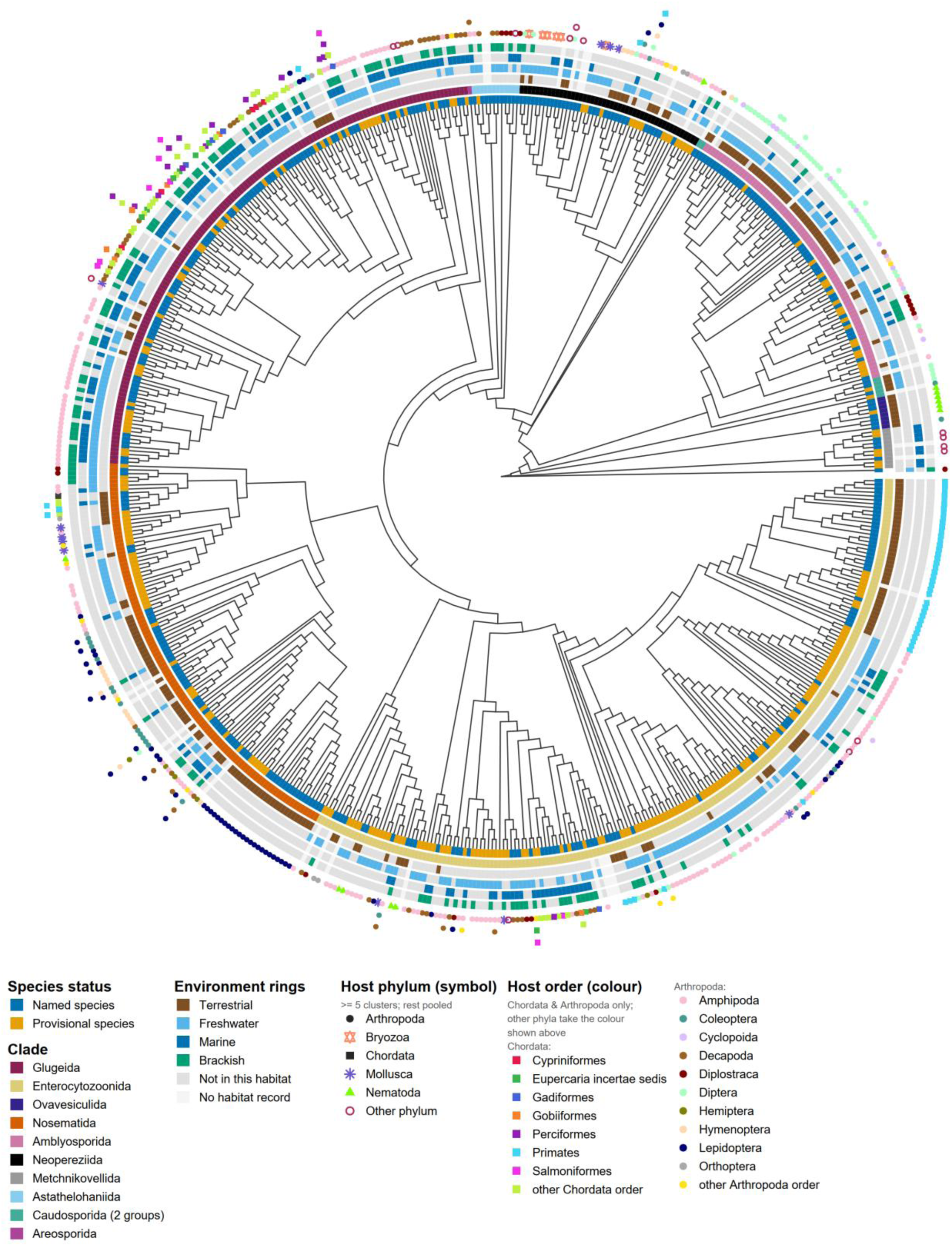
Host range and habitat of microsporidian species-level 18S clusters mapped onto their phylogeny. Maximum-likelihood phylogeny of 609 microsporidian 18S rRNA clusters, each representing sequences grouped at 98% identity; only clusters with at least one reported host are shown. Six rings surround the tree. From the tips outward: species status, clade assignment; and presence of the cluster’s hosts in terrestrial, freshwater, marine, and brackish habitats. Symbols outside the rings denote host taxonomy, one symbol per host taxon of that cluster, stacked outward. Symbol shape indicates host phylum; symbol colour indicates host order within Chordata and Arthropoda, and host phylum otherwise. Host phyla and orders recorded from fewer than five clusters are pooled as “Other phylum” and “other Arthropoda or Chordata order”. Host and habitat records combine GenBank metadata for all sequences in a cluster with curated records from the species attribute database; where a species name spans several clusters, curated records were assigned only to the cluster containing that database entry’s accessions. The minor clade Caudosporida is not recovered as monophyletic in this tree; its two constituent groups are shaded separately.

Host taxonomy and habitat show clear phylogenetic structure. Related clusters frequently infect related hosts: for example, *Nematocida* species form a clade of nematode-infecting species, clusters infecting fish are concentrated within Glugeida and Enterocytozoonida, and Amblyosporida is dominated by species infecting freshwater arthropods such as dipteran insects. Consistent with our previous analysis [4], 18S rRNA identity predicts host taxonomy: among cluster pairs with >95% identity, 99% share a host phylum, 94% a class, 85% an order, 75% a family, 68% a genus and 63% a species, compared with 50%, 23%, 10%, 5%, 3% and 3% when host assignments are shuffled among clusters (S5A Fig). A simple nearest-neighbour predictor, which assigns each cluster the hosts of its most similar cluster, recovers the correct host phylum, class, order, family, genus and species for 91%, 83%, 67%, 61%, 52% and 40% of clusters overall, but its accuracy depends strongly on how close that neighbour is. For clusters whose nearest neighbour is >95% identical, family, genus and species are predicted correctly in 75%, 67% and 57% of cases, falling to 50%, 35% and 24% when the nearest neighbour is 85–90% identical (S5B Fig). Habitat likewise tracks the tree, with marine-host clusters concentrated in Glugeida and terrestrial environmental having the highest percentage in Ovavesiculida and Nosematida.

This structure is not absolute, however, as closely related clusters can infect hosts from different phyla and habitats, indicating repeated host switching amongst these parasites. Provisional, sequence-defined clusters are distributed throughout the tree, and in some regions form groups with no closely related named species, revealing lineages known only from sequencing.

The phylogenetic clustering also reveals diversity within described species. Most strikingly, sequences assigned to the human pathogen *Vittaforma corneae* were split across 30 clusters, which together form a large group within Enterocytozoonida interleaved with 9 provisional clusters (Fig 2, S4 Fig). In total, 24 described species were represented by more than one cluster. Mean pairwise 18S identity between clusters of the same species ranged from 86% to 98.5% (Fig 3A). The two species with the most clusters, *V. corneae* (251 sequences, 30 clusters) and *Nosema bombycis* (140 sequences, 19 clusters), contain many cluster pairs with less than 95% identity: half of all *V. corneae* cluster pairs and a third of *N. bombycis* cluster pairs (Fig 3B-C). Together, this level of 18S divergence within described species indicates substantial unrecognized diversity and suggests that several of these named species may contain sequences from more than one species.

**Fig 3.**
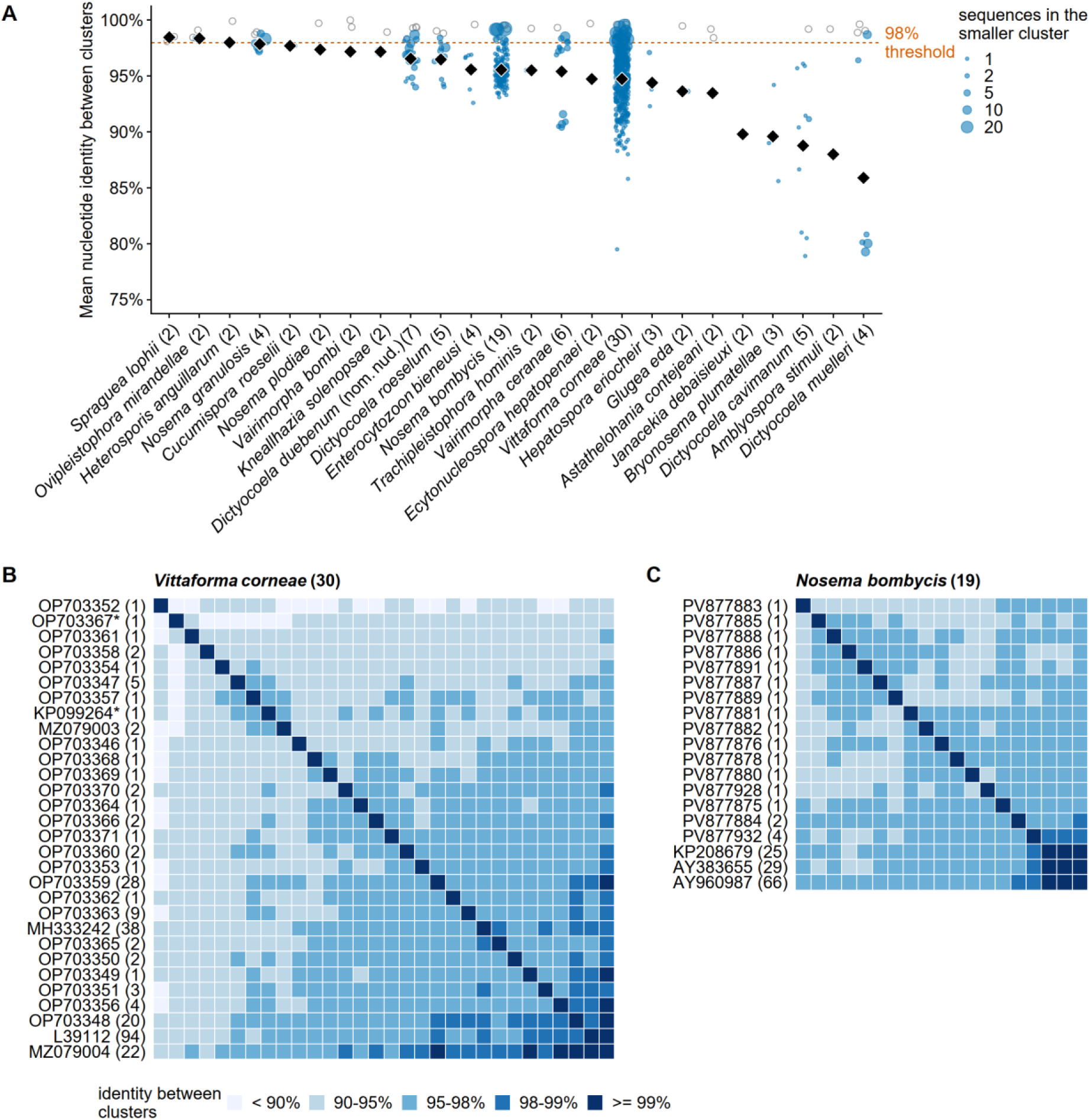
18S rRNA clusters assigned to the same species are often less than 95% identical. **(A)** Identity between clusters for the 24 named species whose sequences were split across more than one cluster at the 98% threshold (116 clusters, 699 cluster pairs). Each blue point is one pair of clusters of the same species, placed at the mean pairwise identity between the sequences of the two clusters and sized by the number of sequences in the smaller cluster of the pair; hollow grey points give the mean identity among the sequences within a cluster, for clusters with at least two sequences; black diamonds give the mean over all cluster pairs of a species. Only sequences of at least 1,000 bp were used, with each cluster’s centroid always included. Species are ordered by their mean between-cluster identity, and the number of clusters is given after each name; the dashed line marks the 98% clustering threshold. **(B, C)** Identity between every pair of clusters of *Vittaforma corneae* (30 clusters, 251 sequences) (B) and *Nosema bombycis* (19 clusters, 140 sequences) (C), coloured by identity class according to the legend at the bottom. Rows and columns share one order, from hierarchical clustering of the identities. Clusters are labelled by the accession of their centroid sequence and, in parentheses, the number of sequences they contain; an asterisk marks a centroid sequence shorter than 1,000 bp.

## Discussion

This work provides the largest database of microsporidia species attributes assembled to date, covering 1,795 species and integrating curated descriptions with sequence-defined diversity from GenBank. These data allow the diversity of the phylum to be described systematically across time, geography, hosts, environments, infected tissues, and spore morphology, and provide a foundation for comparative analyses, such as identifying attributes that correlate with host range or habitat. We show that phylogenetic relatedness is associated with host similarity and can be used to predict the hosts of related lineages, with accuracy that increases with 18S identity; improved modeling approaches, as developed for other parasites, may further enhance this performance [17,18].

Our analysis shows that 98% 18S identity is a useful threshold for delimiting species. A previous analysis of pairwise identity across 53 microsporidia 18S sequences recommended 99% as a useful cutoff [13]. Although a single identity threshold would be convenient for delineating species, no such threshold is likely to be absolute: distinct species can share greater than 99% 18S identity — particularly within the genus *Nosema* — and in some cases greater than 99.9% [19–21]. Conversely, our clustering reveals that considerable diversity remains unappreciated within many described species, in particular *V. corneae*, a cause of human keratitis, and *N. bombycis*, each of which may comprise multiple species — a possibility with practical implications for identifying these pathogens. Diversity within the *Vittaforma* genus has been observed previously, and whole-genome sequencing could help resolve these lineages and determine whether *V. corneae* represents a species complex [16,22–24].

Beyond describing diversity, this database can serve as a living taxonomic reference for microsporidia. New species descriptions can be compared against the recorded attributes and 18S clusters of all existing species to determine whether a putative new species is distinct, and species matching a provisional cluster can then be formally described. The database can be kept current through two complementary routes: formally described species, which researchers can submit through a web portal we have developed (https://www.reinkelab.org/microsporidiaspecies), where attributes are extracted with machine-learning assistance and reviewed before incorporation; and sequence-defined species, which are deposited in GenBank and incorporated by periodic reclustering. As a growing proportion of microsporidia diversity is uncovered through sequencing, both from targeted metagenomic surveys and from incidentally infected hosts, this second route is likely to become the primary way that new species are recognized [15,25,26]. Incorporating community-derived data will keep the database a current reference for describing and organizing the diversity of microsporidia.

## Materials and methods

### Expansion of the microsporidia species attribute database

The database described previously [4] was expanded with species reported since April 2021. Additional species were identified using LLM-assisted literature searches. ChatGPT 5.6 Sol (OpenAI) was used to identify publications describing potentially missing microsporidian species and to compare candidate names, synonyms, and accessions against the database. Only species supported by a complete PDF available online were considered. Candidate additions were independently checked using Claude Opus 4.8 (Anthropic); disagreements were resolved by manually inspecting the source papers, and no species was added solely from LLM output. Attributes were recorded as before, with each row representing a microsporidium and parenthesized text denoting the spore class, host, or measurement condition to which an attribute corresponds. Species were classified as named where the record carried a valid binomial, and as provisional where the name lacked a specific epithet, used a numeric placeholder, or began with an uninformative designation such as “uncultured” or “unnamed”.

### Identification of species from 18S rRNA sequences

Microsporidia 18S rRNA sequences were retrieved from GenBank on August 6, 2026 and trimmed to the region matching the Rfam microsporidia SSU covariance model RF02542 using cmsearch from Infernal v1.1.4 [27]. Separate matches to the model within one sequence were merged when they occurred in the same order along both the sequence and the model, and sequences were retained at an E value of ≤1 × 10⁻⁵. Trimmed sequences were clustered at 98% identity with VSEARCH v2.21.1 [28] using a minimum sequence length of 600 bp and defining identity over the aligned region excluding terminal gaps. Centroid sequences are listed in S1 File. Host, locality, and date (publication year or, if unavailable, the submission year, collection year, or record date) metadata were retrieved for each accession from NCBI Nucleotide using E-utilities [29].

Each cluster was matched to the curated attribute database by shared 18S accession, or, where no accession was shared, by normalised species name. Clusters were matched to the database independently and were never merged with one another; where the clustering split a species across more than one cluster, each cluster is reported separately. For clusters matching an existing entry, the cluster’s hosts, localities, and accessions were recorded in the cluster table (S2 Table) but were not merged into that species’ attribute record, so that the attributes of named species in S1 Table reflect only their published descriptions. For the host taxonomy and environment analyses described below, however, a working copy of the database was generated in which these sequence-derived host records were appended to their matched entries, so that every host shown in the phylogeny could be assigned a taxonomy and environment; S1 Table itself was not modified. Clusters without a database match and with at least one member sequence annotated with a host were added as provisional species with a unique identifier (e.g. MSP-C0042). Where a species name spanned more than one cluster, curated attributes from the database were assigned to a single cluster: the one containing the most of that database entry’s 18S accessions, with ties resolved to the cluster containing the first-listed accession. All other clusters of the same name retained only the host and locality records reported in GenBank for their own member sequences. Clusters with at least one reported host were retained for phylogenetic analysis; S2 Table lists each cluster with its representative accession, phylogenetic clade, all constituent accessions, and its host, locality, and habitat attributes. Each cluster’s host list comprises the hosts reported in GenBank for all of its member sequences, together with the curated hosts of the database record it owns. The attribute analyses (Fig 1) use S1 Table, in which the attributes of named species derive only from their published descriptions, and GenBank-derived information enters only through the provisional cluster species. The phylogenetic and sequence-identity analyses (Fig 2 and 3 and S4 and S5 Figs) use the clusters themselves (S2 Table), which combine the GenBank metadata of their member sequences with the curated attributes of the database record each cluster owns.

### Analysis of microsporidia discovery date and geographical location

For each species, the earliest year in the “Date Identified” column was extracted, and species without year data were excluded. Countries were extracted from the “Locality” column by matching country names, historical country names, and subnational units anywhere in the text, matching longer names before shorter ones so that names could not be matched inside other names. Each species was counted once per country, so a species reported from several countries contributes to each. Species reported from countries that no longer exist were counted as the largest country that occupies those lands (e.g., species from the USSR were counted as Russia). Species reported only from open water were counted for the corresponding ocean or sea and were not assigned to a country. Maps were drawn in R with sf and ggplot2 using Natural Earth 1:50m country boundaries (public domain, via rnaturalearthdata), in Robinson projection [30,31].

### Determination of host taxonomy and habitat

Host names were extracted from both the “Natural Host(s)” and “Experimental Host(s)” columns, with only the natural hosts used for the analysis. Records were separated on semicolons, and on commas only when at least two of the resulting fragments were binomials; parenthesized synonyms were retained as alternative queries, and roles, common names, and subgenera were recorded separately. Abbreviated genus names (e.g., C. tarsalis) were expanded to the genus named in full earlier in the same record where one shared the initial, and otherwise to a genus named elsewhere in the database, the latter only when the expansion was unambiguous — either exactly one genus in the database beginning with that initial was recorded with that specific epithet, or only one genus with that initial occurred at all. Abbreviations consistent with more than one genus were left unexpanded. Where a host record described a hyperparasitic infection, only the direct host of the microsporidium was counted, and the host of that host was recorded separately; for example, in “*Selenidium pygospionis* (direct host); *Pygospio elegans* (polychaete hyperhost),” only *S. pygospionis* was counted as a host.

The taxonomy of each host was determined by querying GBIF (https://www.gbif.org/), the Catalogue of Life (https://www.catalogueoflife.org/), WoRMS (https://www.marinespecies.org/), and NCBI Taxonomy (https://www.ncbi.nlm.nih.gov/taxonomy). A result was discarded if the returned genus differed from the query by more than one character, if the returned specific epithet differed in stem, or if the lineage was placed in a kingdom that microsporidia do not infect. Corrections of two characters were accepted only when a second database independently returned the same corrected name. When databases returned different phyla, the phylum was assigned by majority, after reconciling names that differ between databases (Myzozoa and Miozoa were treated as Apicomplexa, and Discosea and Tubulinea as Amoebozoa); hosts for which no majority existed were left unresolved. The host taxonomy was manually inspected and in two cases, hosts categorized as Echinodermata or Bacteroidota were determined to be incorrect and discarded. The taxonomy assignments for each host are in S3 Table.

Host environment was determined from the host organism only, using WoRMS environment flags, GBIF habitat records, and the Catalogue of Life, defining environment using the union of all the hosts. Hosts for which none of the taxonomic databases returned a habitat were assigned using a table mapping 1,606 host genera to habitats (S4 Table). The table was generated by a large language model (Claude Opus 5, Anthropic) prompted to assign each genus in our host list to one or more of the four habitat categories. Each genus carries a primary habitat, the full set of habitats it occupies, and a confidence level; genera with aquatic larvae and terrestrial adults were assigned both freshwater and terrestrial. The environment of each microsporidia species was taken as the union of the environments of all of its hosts. The habitat assignments for each host are in S3 Table.

### Categorizing and counting tissues infected

Infected tissue names were extracted from the “Site of Infection” field and sorted into 17 categories. Where a record named several tissues, the first tissue keyword in each term determined the category, so that positional descriptors such as “subcutaneous” or “under the peritoneum of” were assigned to the tissue they modify. Each species was counted once per category, so a species infecting several tissues contributes to each. Records describing systemic or generalized infection were assigned to a “Systemic / general” category and retained; where a species reported both a systemic and a specific site, the specific site was also counted.

### Spore shape classification and spore morphology analysis

Spore shapes were extracted from the “Spore Shape” column. Each descriptor was assigned to one of six classes (elliptical, oval, ovocylindrical, pyriform, rod-shaped, spherical) by matching against an ordered list of terms per class, with more specific classes tested first so that compound descriptors were not assigned to the more general class. Descriptors naming a shape outside these classes were classified as “Other”. Where an entry reported several descriptors, those from fresh or live preparations were used in preference to fixed or stained ones, and those of ordinary spores in preference to macrospores, meiospores, or aberrant spores.

Spore lengths and widths were extracted from the “Spore Length Average” and “Spore Width Average” columns, using the same preparation priority as for shape. Stated means were used in preference to the midpoints of reported ranges, and length and width were taken from measurements of the same spore class; entries where they could not be matched were excluded. Polar tubule coil numbers were taken from the reported average, or from the midpoint of the reported range where only a range was given.

### Phylogenetic analysis

Centroids of microsporidia 18S sequences (*N* = 680) obtained as described above were aligned with MUSCLE v5.1 [32], and columns present in fewer than 10% of sequences were removed with trimAl v1.5 (-gt 0.1) [33], yielding an alignment of 1,403 columns. The best-fitting substitution model was selected with ModelFinder under the Bayesian information criterion (-m MFP), which returned GTR+F+R10. A maximum-likelihood phylogeny was inferred with IQ-TREE v3.0.1 [34] under this model with 1,000 ultrafast bootstrap replicates, 1,000 SH-aLRT replicates, and NNI optimisation to reduce model-violation bias (-B 1000 --alrt 1000 --bnni), and rooted on *Mitosporidium daphniae* (MF278562.1). The tree was pruned to only include sequence clusters that were annotated with at least one host, resulting in 609 tips. The clade assignment of each species was compared to a previous assessment, and all overlapping species shared the same assignment between the two studies [3].

### Relationship between 18S rRNA identity and host taxonomy

All sequences of at least 1,000 bp (2,277 sequences from 411 clusters) were compared with VSEARCH (--allpairs_global), defining identity over the aligned region excluding terminal gaps as in the clustering [28]; clusters with no sequence of this length were excluded. Identity between two clusters was defined as the mean of the pairwise identities between all retained sequences of one cluster and all retained sequences of the other. Clusters were not merged by species name, and each cluster carried the host list given in S2 Table, with host taxonomy determined as described above. Two clusters were scored as sharing a taxon at a rank if any host of one belonged to the same taxon as any host of the other. For nearest-neighbour prediction, each cluster was assigned the host list of the cluster with the highest identity to it, and the prediction was scored as correct at a rank if the two clusters shared a taxon. Both quantities were summarised in 5% identity bins. A null distribution was generated by shuffling host lists among clusters 10,000 times with the identity matrix held fixed. Each bin was compared with the same bin of the shuffled data by one-sided permutation tests in both directions, p = (k + 1)/(N + 1) with k the number of shuffles at least as extreme as the observed value, and p values were adjusted across the bins and ranks of each panel by the Benjamini–Hochberg procedure.

### Sequence identity between clusters assigned to the same species

For each named species represented by more than one cluster, pairwise identities between all of its sequences were computed with VSEARCH (--allpairs_global), defining identity over the aligned region excluding terminal gaps as in the clustering [28]. Sequences shorter than 1,000 bp were excluded from the identity calculations, except that the centroid of every cluster was retained regardless of length so that no cluster was lost (718 of 1,962 sequences retained across 24 species and 116 clusters). Identity between two clusters was defined as the mean of the pairwise identities between all retained sequences of one cluster and all retained sequences of the other, and identity within a cluster as the mean among its retained sequences; the value for a species is the mean over all pairs of its clusters.

## Figure generation

All figures were generated in R version 4.6.1 [35] using ggplot2 [30] and assembled with patchwork [36].

## Acknowledgements

This work was supported by a Canadian Institutes of Health Research grant (no. 461807 to A. W. R.).

## Data availability

All microsporidia species, sequences of 18S centroids, and host data are available in S1 File and S1–S4 Tables. The most recent version of the microsporidia species data can be found at https://www.reinkelab.org/microsporidiaspecies. All code used for data curation, analysis, and figure generation is available at https://github.com/reinkelab-utoronto/microsporidia-species-database.

## Declaration of generative AI and AI-assisted technologies in the manuscript preparation process

During the preparation of this work the authors used Claude Opus 4.8 and 5, Claude Fable 5.1, ChatGPT 5.6 Sol, and ChatGPT 6 Astra to generate Python scripts for processing the data, R scripts for creating figures and tables, to generate draft text for the paper, and to identify and extract species attribute information from papers reporting novel microsporidia species. The authors reviewed and edited the content and take full responsibility for the content of the published article.

## Author contributions

A. W. R. conceptualized the study. C. C. and A. A. curated microsporidia species information and generated draft code for analysis. A. W. R. designed and performed all analyses in the paper. The paper was written by A. W. R. and was approved by all authors. Mentorship and funding acquisition were provided by A. W. R.

## Supporting information

**S1 File. FASTA file of the trimmed centroid 18S rRNA sequences for each microsporidia cluster with at least one reported host.**

**S1 Table. Microsporidia species attribute database.**

**S2 Table. Microsporidia 18S rRNA sequence clusters and their clade, status, accessions, hosts, countries, and associated curated database record.**

**S3 Table. Host taxonomy and habitat assignments for each host, combining the taxonomic databases and the genus-level habitat predictions.**

**S4 Table. Host habitat predictions from the LLM-generated genus-level habitat table.**

## Supporting figures and figure legends

**S1 Fig.**
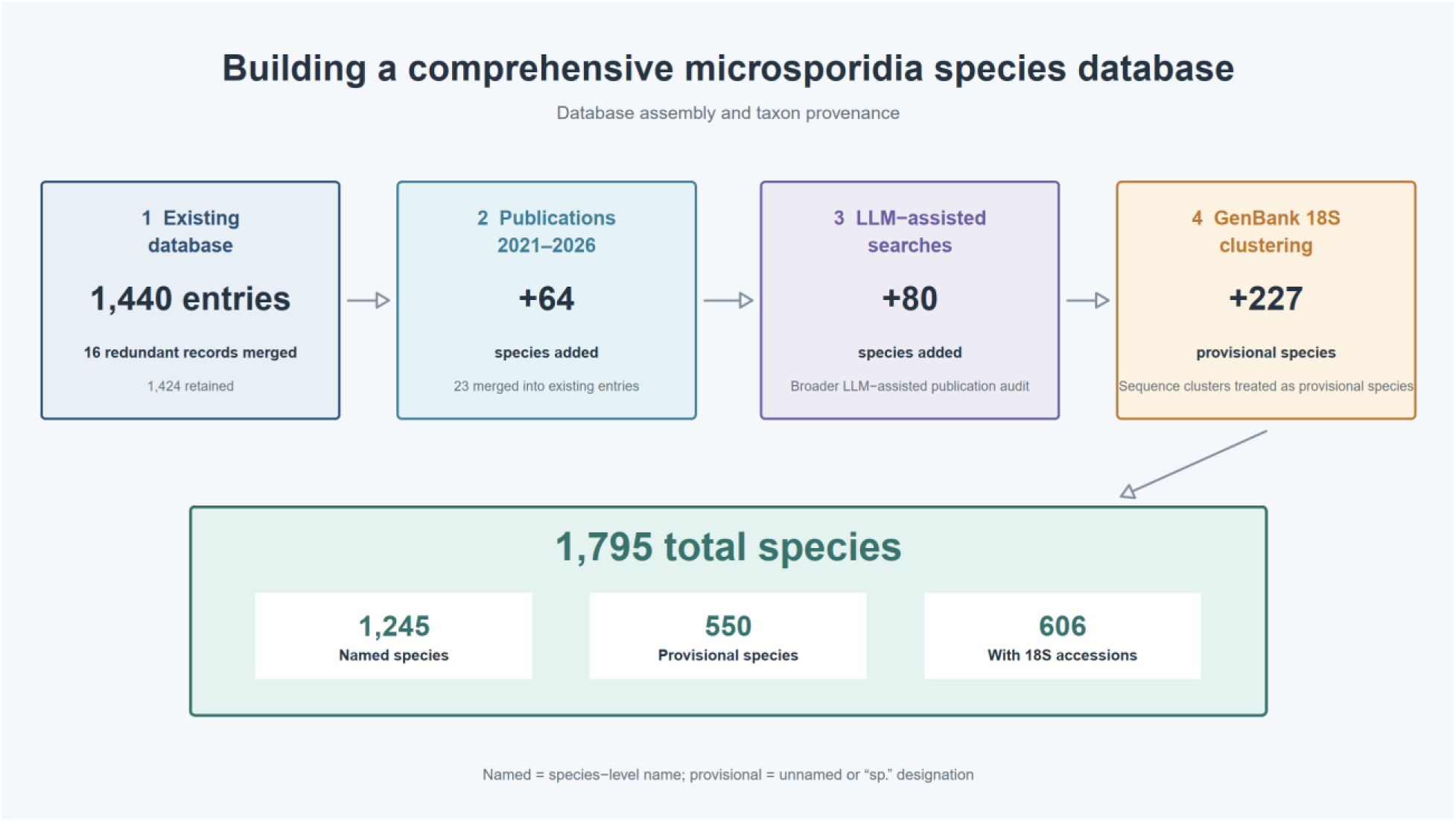
Expansion and curation of the microsporidia species attribute database. The existing database contained 1,440 entries. Consolidation of 16 redundant records resulted in 1,424 retained species records. Species described from 2021 to 2026 were identified from supplied publications and targeted literature searches, adding 64 species not already represented in the database as well as updating 23 existing entries. Broader LLM-assisted searches of the literature identified 80 additional species. Clustering of GenBank 18S rRNA sequences added 227 sequence-defined provisional species. The final database contains 1,795 species, comprising 1,245 named species and 550 provisional species, with 606 species represented by at least one 18S rRNA accession. Named species were defined as records with a species-level name, whereas provisional species lacked a species name or were designated “sp.”

**S2 Fig.**
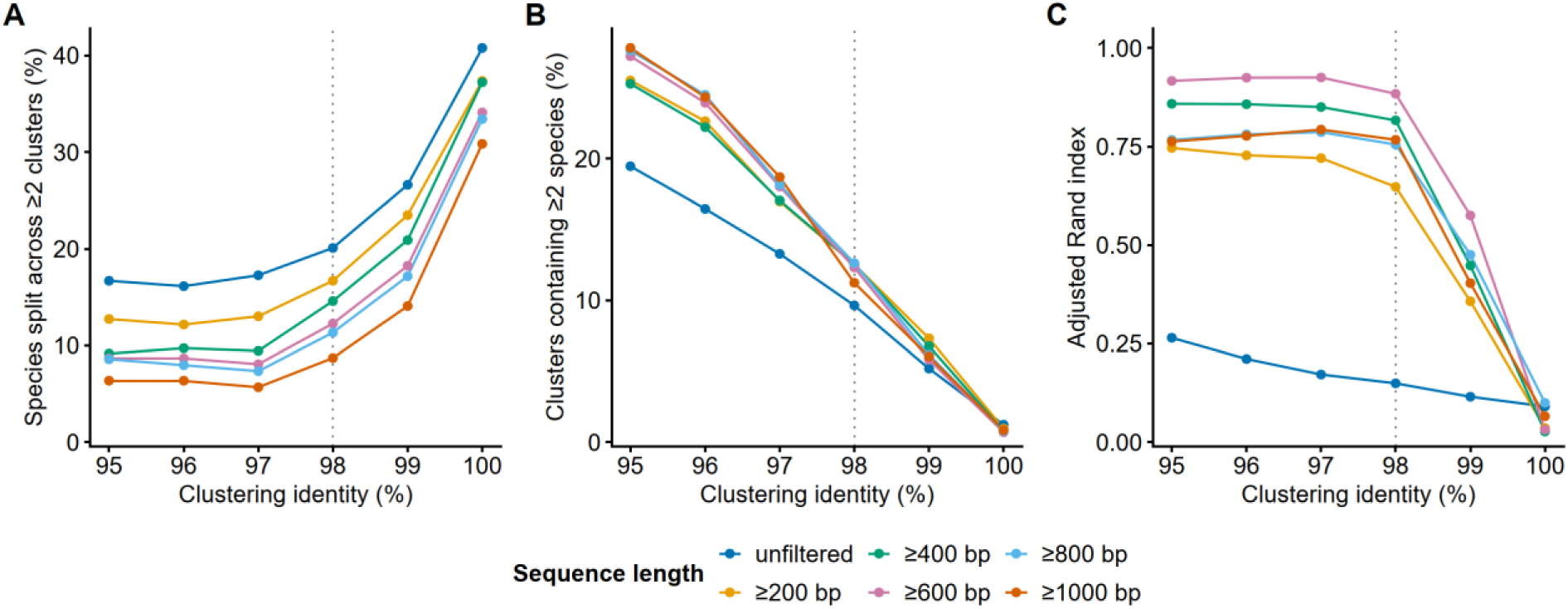
Selection of clustering thresholds for microsporidia 18S rRNA sequences. Trimmed 18S rRNA sequences were clustered with VSEARCH at sequence identity thresholds from 95% to 100% after filtering by trimmed sequence length. The sequence-length threshold is colored according to the legend at the bottom with unfiltered retaining all sequences. The resulting clusters were scored against species binomials from sequence metadata (n = 12,641 sequences from 353 microsporidia species; sequences without species-level names were clustered but not scored). **(A)** Percentage of species whose sequences were assigned to two or more clusters. **(B)** Percentage of clusters containing two or more species. **(C)** Adjusted Rand index measuring overall agreement between clusters and species labels (1, perfect agreement; 0, agreement expected by chance). Dotted lines indicate the chosen threshold of 98% identity.

**S3 Fig.**
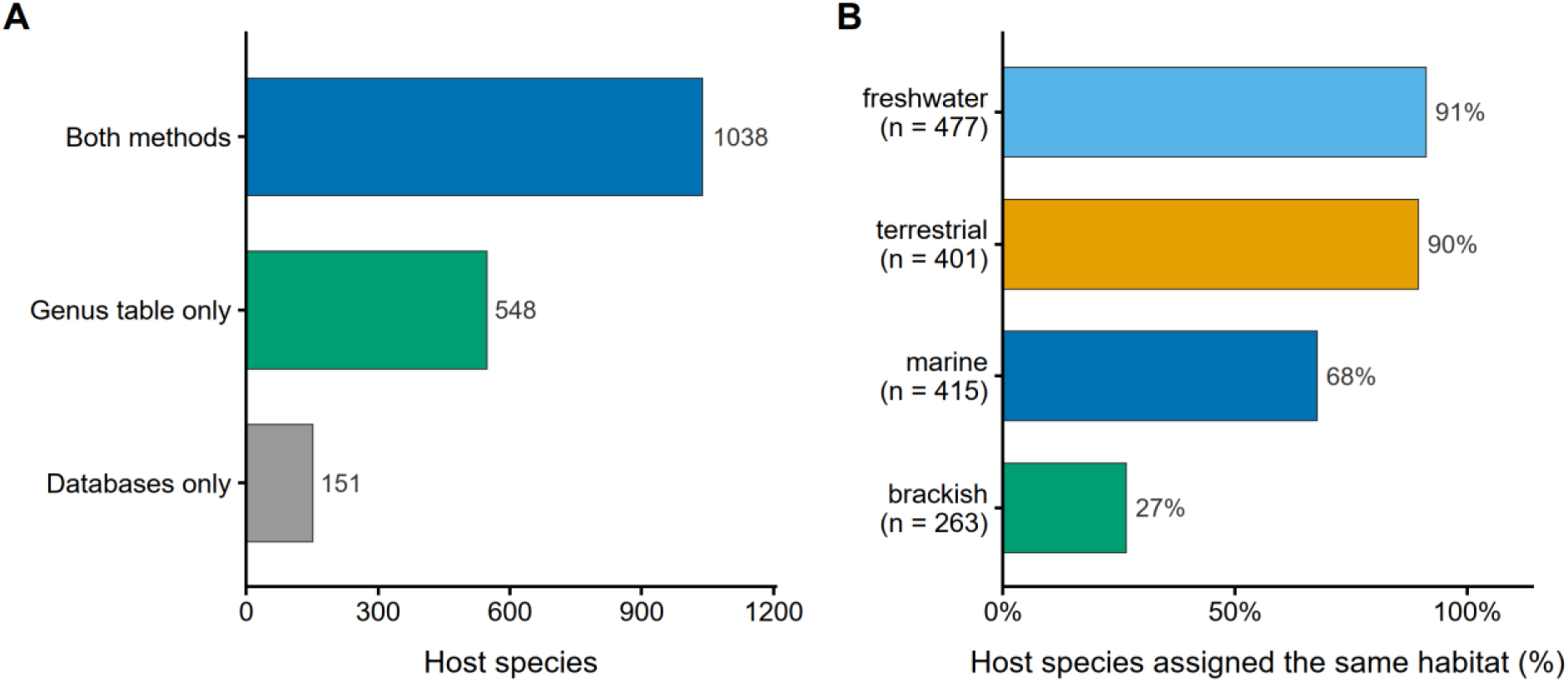
Concordance between database-derived and inferred host habitats. **(A)** Number of host species assigned a habitat by the taxonomic databases (GBIF, WoRMS, Catalogue of Life), by the genus-level habitat table, or by both. **(B)** For host species assigned a habitat by both, the percentage assigned the same habitat by the genus-level table, shown separately for each habitat; *n* under each habitat label gives the number of host species.

**S4 Fig.**
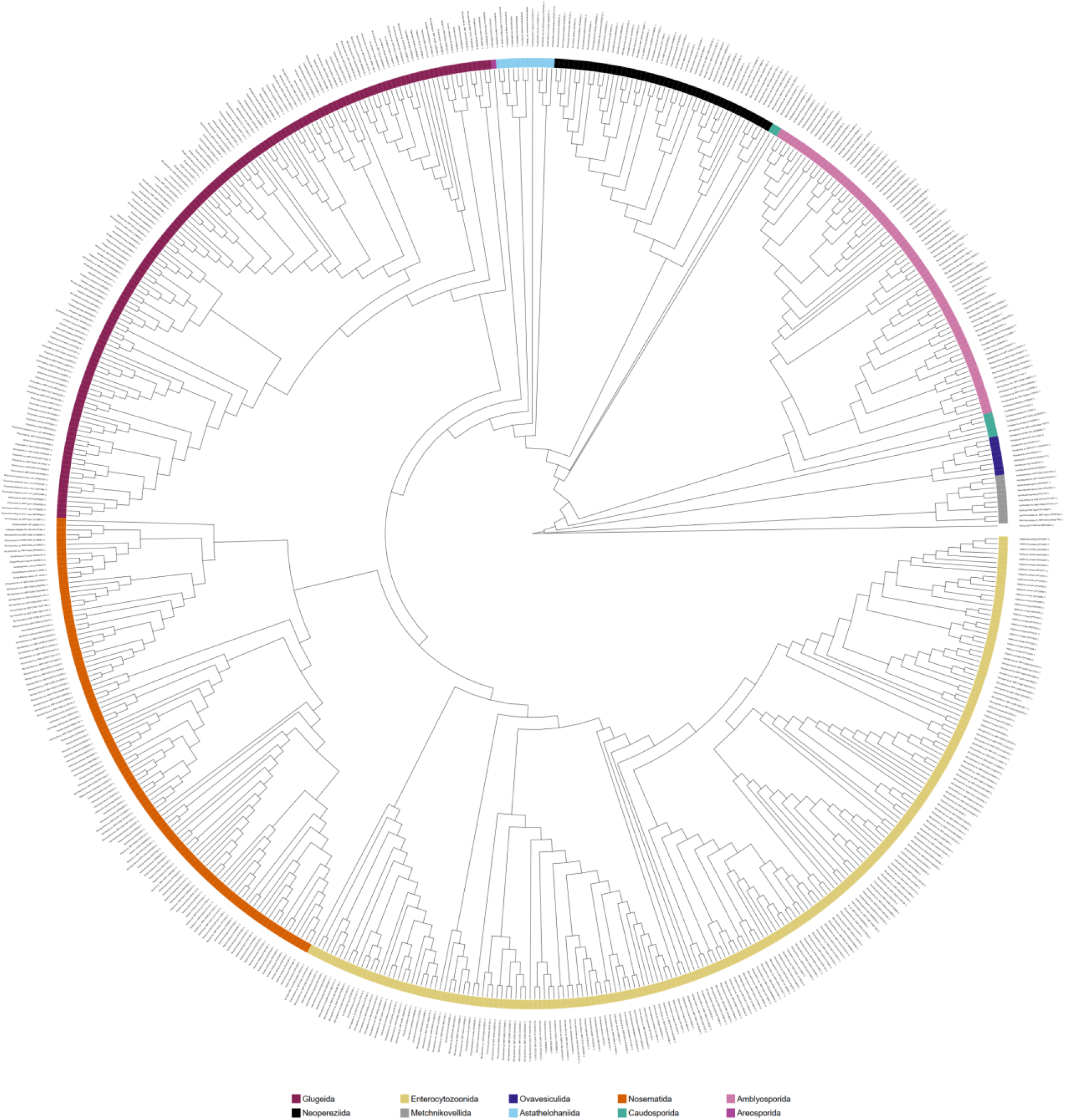
Microsporidian 18S rRNA phylogeny with cluster identities labelled. The tree from Fig 2, with each tip labelled by its species name and the accession of the cluster’s representative sequence. Clusters carrying a valid binomial are labelled with that name; those defined only by sequence and host records are given a provisional identifier of the form “*Genus* sp. MSP-Cnnnn”. Ring colors indicate clade assignment, as in Fig 2. Branch lengths are not shown, and host, habitat and taxonomic annotations are omitted for legibility.

**S5 Fig.**
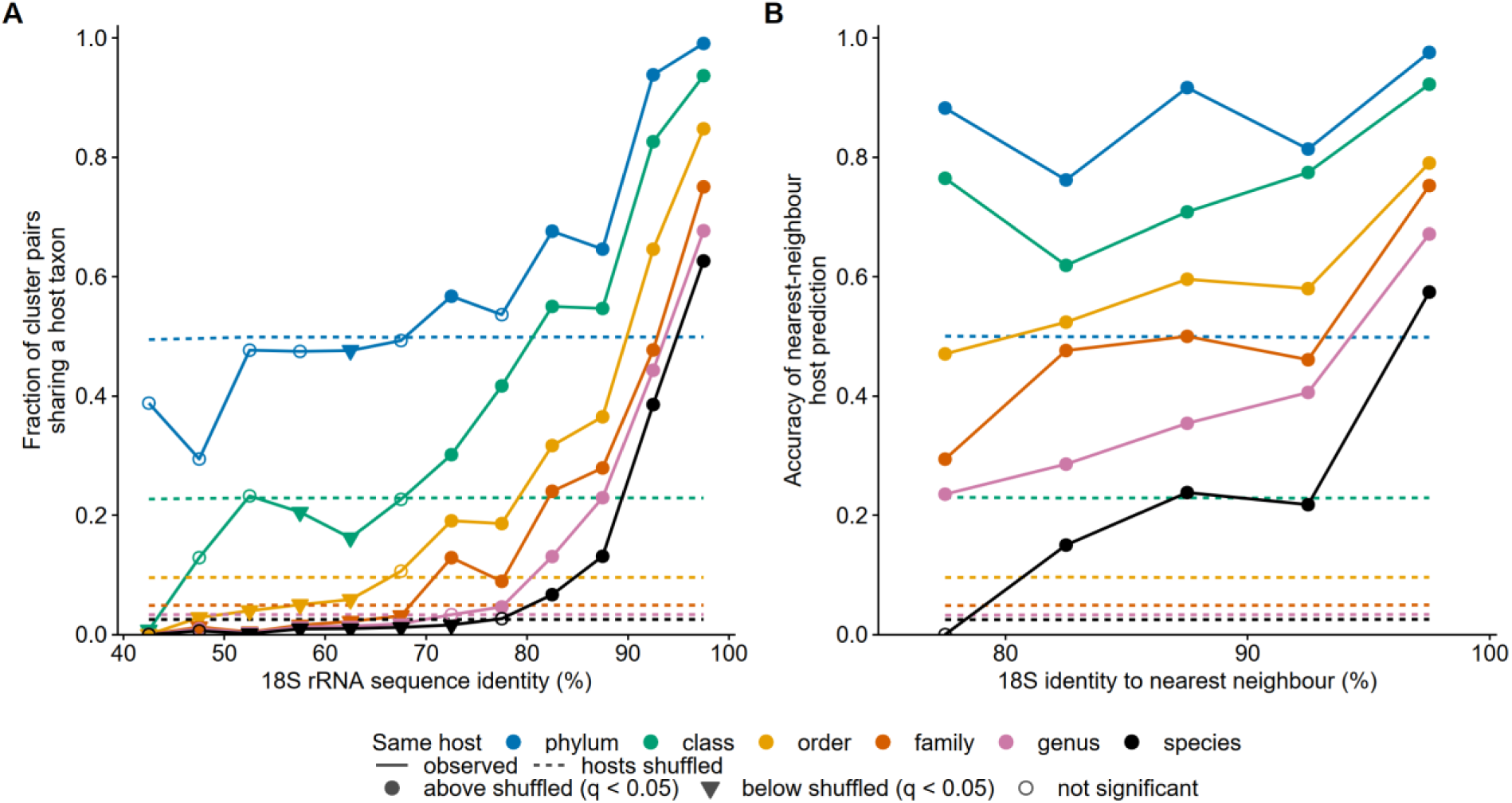
18S rRNA identity predicts host taxonomy. **(A)** Fraction of cluster pairs (n = 84,255 pairs among 411 clusters) that share at least one host taxon at each taxonomic rank, in 5% bins of mean 18S rRNA identity between the clusters. **(B)** Accuracy of a nearest-neighbour predictor that assigns each cluster the hosts of the cluster with the highest 18S identity to it, scored as correct when the two share a host taxon at the given rank, in 5% bins of identity to the nearest neighbour. In both panels solid lines are the observed data and dashed lines of the same colour are the mean of 10,000 shuffles of host lists among clusters with the identity matrix held fixed. Each point was compared with the same bin of the shuffled data by one-sided permutation tests: filled circles lie significantly above the shuffled value, filled triangles significantly below, and open circles are not significant.

## References

1. Vavra J, Lukes J. Microsporidia and “the art of living together”. Adv Parasitol. 2013;82: 253–319. doi:10.1016/B978-0-12-407706-5.00004-6

2. Wadi L, Reinke AW. Evolution of microsporidia: An extremely successful group of eukaryotic intracellular parasites. PLoS Pathog. 2020;16: e1008276. doi:10.1371/journal.ppat.1008276

3. Bojko J, Reinke AW, Stentiford GD, Williams B, Rogers MSJ, Bass D. Microsporidia: a new taxonomic, evolutionary, and ecological synthesis. Trends Parasitol. 2022;38: 642–659. doi:10.1016/j.pt.2022.05.007

4. Murareanu BM, Sukhdeo R, Qu R, Jiang J, Reinke AW. Generation of a Microsporidia Species Attribute Database and Analysis of the Extensive Ecological and Phenotypic Diversity of Microsporidia. mBio. 2021;12: 10.1128/mbio.01490-21. doi:10.1128/mbio.01490-21

5. Han B, Takvorian PM, Weiss LM. The Function and Structure of the Microsporidia Polar Tube. In: Weiss LM, Reinke AW, editors. Microsporidia: Current Advances in Biology. Cham: Springer International Publishing; 2022. pp. 179–213. doi:10.1007/978-3-030-93306-7_8

6. Han B, Pan G, Weiss LM. Microsporidiosis in Humans. Clin Microbiol Rev. 2021;34: e00010-20. doi:10.1128/CMR.00010-20

7. Huang Q, Hu W, Meng X, Chen J, Pan G. Nosema bombycis: A remarkable unicellular parasite infecting insects. J Eukaryot Microbiol. 2024;71: e13045. doi:10.1111/jeu.13045

8. Chaijarasphong T, Munkongwongsiri N, Stentiford GD, Aldama-Cano DJ, Thansa K, Flegel TW, et al. The shrimp microsporidian *Enterocytozoon hepatopenaei* (EHP): Biology, pathology, diagnostics and control. J Invertebr Pathol. 2021;186: 107458. doi:10.1016/j.jip.2020.107458

9. Martín-Hernández R, Bartolomé C, Chejanovsky N, Conte YL, Dalmon A, Dussaubat C, et al. Nosema ceranae in Apis mellifera: a 12 years postdetection perspective. Environ Microbiol. 2018;20: 1302–1329. 10.1111/1462-2920.14103

10. Herren JK, Mbaisi L, Mararo E, Makhulu EE, Mobegi VA, Butungi H, et al. A microsporidian impairs Plasmodium falciparum transmission in Anopheles arabiensis mosquitoes. Nat Commun. 2020;11: 1–10. doi:10.1038/s41467-020-16121-y

11. LeBrun EG, Jones M, Plowes RM, Gilbert LE. Pathogen-mediated natural and manipulated population collapse in an invasive social insect. Proc Natl Acad Sci. 2022;119: e2114558119. doi:10.1073/pnas.2114558119

12. Sprague V. Annotated List of Species of Microsporidia. In: Bulla LA, Cheng TC, editors. Comparative Pathobiology: Volume 2 Systematics of the Microsporidia. Boston, MA: Springer US; 1977. pp. 31–334. doi:10.1007/978-1-4613-4205-2_2

13. de Albuquerque NRM, Haag KL. Using average nucleotide identity (ANI) to evaluate microsporidia species boundaries based on their genetic relatedness. J Eukaryot Microbiol. 2023;70: e12944. doi:10.1111/jeu.12944

14. Shreenidhi PM, Reinke AW. 25 Years Since the First Microsporidian Genome Assembly, Insights on How Genomics Has Shaped Our Understanding of Parasites with the Smallest Eukaryotic Genomes. Annu Rev Microbiol. 2026 [cited 21 Sept 2026]. doi:10.1146/annurev-micro-042424-034850

15. Trzebny A, Slodkowicz-Kowalska A, Becnel JJ, Sanscrainte N, Dabert M. A new method of metabarcoding Microsporidia and their hosts reveals high levels of microsporidian infections in mosquitoes (Culicidae). Mol Ecol Resour. 2020;20: 1486–1504. doi:10.1111/1755-0998.13205

16. Combes L, Chauvet M, Monjot A, Moné A-I, Pont C, Moreau C, et al. Unraveling Long-Term Microsporidia Diversity and Dynamics in Lake Aydat (France) Through Paleogenomics. Environ DNA. 2026;8: e70305. doi:10.1002/edn3.70305

17. Carbajo Jr. AL, Vensko TA, Pellett PE. Sequence based virus host prediction: a curated dataset and generalizable framework for training artificial intelligence to identify viruses of humans. Virus Evol. 2026;12: veag009. doi:10.1093/ve/veag009

18. Klimov PB, He Q. Predicting host range expansion in parasitic mites using a global mammalian-acarine dataset. Nat Commun. 2024;15: 1–14. doi:10.1038/s41467-024-49515-3

19. Reinke AW, Balla KM, Bennett EJ, Troemel ER. Identification of microsporidia host-exposed proteins reveals a repertoire of rapidly evolving proteins. Nat Commun. 2017;8: 14023. doi:10.1038/ncomms14023

20. Kyei-Poku G, Gauthier D, Van Frankenhuyzen K. Molecular Data and Phylogeny of Nosema Infecting Lepidopteran Forest Defoliators in the Genera Choristoneura and Malacosoma. J Eukaryot Microbiol. 2008;55: 51–58. doi:10.1111/j.1550-7408.2007.00302.x

21. Tokarev YS, Huang W-F, Solter LF, Malysh JM, Becnel JJ, Vossbrinck CR. A formal redefinition of the genera *Nosema* and *Vairimorpha* (Microsporidia: Nosematidae) and reassignment of species based on molecular phylogenetics. J Invertebr Pathol. 2020;169: 107279. doi:10.1016/j.jip.2019.107279

22. Chen J-S, Hsu T-K, Hsu B-M, Chao S-C, Huang T-Y, Ji D-D, et al. Swimming Pool–Associated Vittaforma-Like Microsporidia Linked to Microsporidial Keratoconjunctivitis Outbreak, Taiwan - Volume 25, Number 11—November 2019 - Emerging Infectious Diseases journal - CDC. [cited 21 Sept 2026]. doi:10.3201/eid2511.181483

23. Chen J-S, Hsu B-M, Tsai H-C, Chen Y-P, Huang T-Y, Li K-Y, et al. Molecular surveillance of Vittaforma-like microsporidia by a small-volume procedure in drinking water source in Taiwan: evidence for diverse and emergent pathogens. Environ Sci Pollut Res. 2018;25: 18823–18837. doi:10.1007/s11356-018-2081-4

24. Motro Y, Wajnsztajn D, Michael-Gayego A, Mathur S, Marano RB, Salah I, et al. Metagenomic sequencing for investigation of a national keratoconjunctivitis outbreak, Israel, 2022. Eurosurveillance. 2023;28: 2300010. doi:10.2807/1560-7917.ES.2023.28.31.2300010

25. Khalaf A, Zhou C, Weber CC, Vancaester E, Sims Y, Makunin A, et al. Forty new genomes shed light on sexual reproduction and the origin of tetraploidy in Microsporidia. PLOS Biol. 2025;23: e3003446. doi:10.1371/journal.pbio.3003446

26. Lisnerová M, Nayak S, Beest GS van, Bürgerová M, Lövy A, Pecková H, et al. Microsporidian diversity and host associations in aquatic invertebrate communities of South Bohemian fish ponds revealed by metabarcoding. Metabarcoding Metagenomics. 2026;10: e201781. doi:10.3897/mbmg.10.201781

27. Nawrocki EP, Eddy SR. Infernal 1.1: 100-fold faster RNA homology searches. Bioinformatics. 2013;29: 2933–2935. doi:10.1093/bioinformatics/btt509

28. Rognes T, Flouri T, Nichols B, Quince C, Mahé F. VSEARCH: a versatile open source tool for metagenomics. PeerJ. 2016;4: e2584. doi:10.7717/peerj.2584

29. Sayers EW, Bolton EE, Fine AM, Kelly C, Kim S, Landrum M, et al. Database resources of the National Center for Biotechnology Information in 2026. Nucleic Acids Res. 2026;54: D20–D27. doi:10.1093/nar/gkaf1060

30. Wickham H. ggplot2: Elegant Graphics for Data Analysis. Springer-Verlag New York; 2016. Available: https://ggplot2.tidyverse.org

31. Pebesma E. Simple Features for R: Standardized Support for Spatial Vector Data. R J. 2018;10: 439–446. doi:10.32614/RJ-2018-009

32. Edgar RC. Muscle5: High-accuracy alignment ensembles enable unbiased assessments of sequence homology and phylogeny. Nat Commun. 2022;13: 6968. doi:10.1038/s41467-022-34630-w

33. Capella-Gutiérrez S, Silla-Martínez JM, Gabaldón T. trimAl: a tool for automated alignment trimming in large-scale phylogenetic analyses. Bioinformatics. 2009;25: 1972–1973. doi:10.1093/bioinformatics/btp348

34. Wong TKF, Ly-Trong N, Ren H, Demotte P, Baños H, Roger AJ, et al. IQ-TREE 3: phylogenomic inference software using complex evolutionary models. Mol Biol Evol. 2026;43: msag117. doi:10.1093/molbev/msag117

35. R Core Team. R: A Language and Environment for Statistical Computing. Vienna, Austria: R Foundation for Statistical Computing; 2026. doi:10.32614/R.manuals

36. Pedersen TL. patchwork: The Composer of Plots. 2025. doi:10.32614/CRAN.package.patchwork

